# Structural and Dynamical Properties of Elastin-Like Peptides near their Lower Critical Solution Temperature

**DOI:** 10.1101/2022.09.25.509363

**Authors:** Tatiana I. Morozova, Nicolás A. García, Olga Matsarskaia, Felix Roosen-Runge, Jean-Louis Barrat

## Abstract

Elastin-like peptides (ELPs) are artificially derived intrinsically disordered proteins (IDPs) mimicking the hydrophobic repeat unit in the protein elastin. ELPs are characterized by a lower critical solution temperature (LCST) in aqueous media. Here, we investigate the sequence GVG(VPGVG)_3_ over a wide range of temperatures (below, around, and above the LCST) and peptide concentrations employing all-atom molecular dynamics simulations, where we focus on the role of intra- and inter-peptide interactions. We begin by investigating the structural properties of a single peptide that demonstrates a hydrophobic collapse with temperature, albeit moderate, as the sequence length is short. We observe a change in the interaction between two peptides from repulsive to attractive with temperature by evaluating the potential of mean force, indicating an LCST-like behaviour. Next, we explore dynamical and structural properties of peptides in multi-chain systems. We report the formation of dynamical aggregates with coil-like conformation, in which Val central residues play an important role. Moreover, the lifetime of contacts between chains strongly depends on the temperature and can be described by a power-law decay that is consistent with the LCST-like behaviour. Finally, the peptide translational and internal motion are slowed down by an increase in the peptide concentration and temperature.

## Introduction

Elastin is a key protein of the extracellular matrix that provides elasticity to biological tissues such as lungs, arteries, skin, tendons, ligaments, and cartilage with extraordinary long-term stability and resilience.^1^ These properties originate from the nature of its monomer - tropoelastin. The latter is a soluble protein composed of cross-linking and hydrophobic domains.^2^ While the cross-linking domains are believed to be responsible for cohesiveness and durability, the hydrophobic domains increase the ability of tropoelastin to self-assemble and recoil.^3^

Elastin-like peptides (ELPs) are synthetic peptides that mimic these hydrophobic domains.^1,4^ Their sequence is composed of pentapeptide repeat units Val-Pro-Gly-Xaa-Gly (VPGXG), where the guest residue X can be any amino acid except for proline. Apart from their model role to understand elastin’s elasticity, ELPs show multiple stimuli-responsive properties^4^ and are biodegradable.^5^ This makes them highly suitable for a broad range of applications^6^ including advanced biomaterials,^7^ tissue engineering, ^8^ protein purification,^9^ and drug delivery.^4,7,10^ In particular, ELPs typically exhibit a lower critical solution temperature (LCST) phase behavior in aqueous environments *in vitro* ^4^ where initially coiled-like chains undergo a conformation transition into globule-like upon increasing the temperature. The transition temperature *T*_t_ at which this conformation transition occurs depends on various parameters such as the peptide sequence and length^11–13^ and can be regulated by external stimuli such as pH,^14^ ion concentration,^15^ and pressure.^16^ In addition, ELPs are good models for intrinsically disordered proteins which have a profound impact on cellular processes and pathologies^17^ and thus require systematic physicochemical understanding.^18–20^

The origin of the LCST-like phase behavior of ELPs has been examined by a variety of experimental techniques including microscopy,^21^ differential scanning calorimetry, ^22^ dielectric relaxation,^23^ scattering^24^ and different spectroscopic techniques.^25,26^ In general, the LCST-like phase behavior has been linked to hydrophobic effects that increase with temperature and the formation of secondary structures, in particular *β*-turns. Urry et al^27^ described this transition as a multi-step process that involves a conformational transition of individual chains from the coil to the so-called *β*-spiral model (the secondary structure is rich in *β*-turns) in which hydrophobic groups are exposed to the solvent that drives interchain aggregation via hydrophobic interactions. In contrast, more recent experimental work that includes NMR^28,29^ and CD spectroscopy^30^ indicated that the secondary structure of ELPs is rich in *β*-turns also below the LCST. Thus, the controversy about the structural properties of ELPs in the LCST temperature range remains. This is not surprising since conformational heterogeneity and self-aggregation of ELPs make their experimental characterization challenging.

Computer simulations can provide detailed information on the structural and dynamical properties of ELPs near LCST. However, until quite recently a large body of work was dedicated to single-chain systems due to computational limitations. Tarakanova et al^31^ demonstrated that peptide sequence, chain length and salt concentration influence the conformation transition at a single-chain level using 3-6 pentamers sequence. Zhao et al^13^ investigated a single (VPGVG)_*n*_ peptide for various chain lengths *n*. Their work revealed a power-law dependence of the transition temperature *T*_t_ on the chain length, however, the propensity of hydrogen bond formation depends solely on a pentamer sequence. Additionally, it was pointed out that short ELPs such as those investigated here do not exhibit conformational transition as their transition temperature lays above the boiling point of water. Zhao and Kremer^32^ demonstrated that the conformational transition of a (VPGVG)_30_ polypeptide strongly depends on proline isomerisation.

Only a few studies have addressed ELPs properties when partitioning into aggregates from all-atom molecular dynamics (MD) simulations. Li et al^33^ investigated the early stage of aggregation by considering a dimer of VPGVG_18_, where structural properties of the chains resemble those of single-chain systems. The authors proposed that moderate changes at a single-chain level occurring over *T*_t_ are sufficient to drive peptide aggregation. Next, Rauscher and Pomes^34^ considered an assembly of 27 short ELPs with the sequence GVPGV_7_. Their study suggests that conformational ensembles visited by a single chain in solution or in an aggregate differ significantly. Even though the peptide aggregation is partly driven by hydrophobic effects, the self-assembly does not result in the formation of a hydrophobic core, the peptide backbone remained hydrated. In the most recent work by Li et al^35^ the influence of the pentamer sequence, namely VPGVG and VGPVG, on the aggregation behavior of 27 long ELPs with 18 pentameric units was considered. In agreement with an earlier study, ^34^ the authors concluded that for these sequences no hydrophobic core was formed and the individual chain conformation is more extended when chains partition into a cluster. The difference in the sequence resulted in higher hydrophobicity of poly(VPGVG) aggregates with fewer intrachain interactions and stronger interchain hydrogen bonds compared to poly(VGPVG).

In this work, we focus on the phase behavior of the sequence GVG(VPGVG)_3_ as a function of temperature and the peptide concentration. This sequence has been extensively studied in experiments since its synthesis could be performed by solid-phase peptide synthesis, and could provide a comprehensive few on the role of intra- and inter-peptide interactions at LCST. Additionally, most computational studies of ELPs employed the force fields that rely on the TIP3P water model for modeling the solvent. However, this water model describes the structural and dynamical properties of real water reasonably well only at room temperature while being inaccurate at other temperatures. ^36^ We believe that adequate modeling of the solvent is essential in studying the LCST-like behavior of the ELPs since interactions between solute and solvent play a key role in the conformational transition. Thus, we employ allatom MD simulation in conjunction with state-of-the-art force fields shown to be suitable for modeling IDPs and advanced sampling methods to elucidate how structural and dynamical properties of a peptide and its aggregates might be affected by temperature and crowding effects, i.e. an increase in peptide concentration.

This paper is organized as follows. First, we describe the numerical model and simulation details. Next, we present the structural properties of a single-peptide system near temperatures where the LCST-like behavior is expected. Then, we consider the interaction between two peptides at the same temperature range through MD simulations combined with the umbrella sampling technique. Finally, we consider multi-peptide systems across a variety of temperatures and peptide concentrations.

## Simulation Model and Methods

We perform explicit solvent, all-atom MD simulations employing the AMBER99SB-ILDN force field^37^ for the peptide and the TIP4P-D water model. ^38^ This choice is motivated by previous works showing that commonly used water models, e.g., TIP3-P, might not fully capture the strength of dispersion interactions in protein-water systems. Thus, the conformational properties of the disordered proteins were too compact in comparison with experimental values.^38,39^ Moreover, Henriques and Skepö demonstrated that when the TIP4P-D water model is used in conjunction with the standard force field AMBER99SB-ILDN for modeling IDPs, the simulations results are in good agreement with the experiments as well as with simulations in which IDP-suitable force fields were employed. ^40^ Hence, we use this combination of the force field and water model in our study. Additionally, it was shown that the TIP4P-D water model reproduces well thermodynamic, dynamic, and dielectric properties of liquid water in a wide temperature range from 280 K to 350 K which is of interest here.^36^

The initial peptide configuration is assembled using the Avogadro molecular builder (ver. 1.20).^41^ In our modeling, we consider neutral pH values which imply that the N− and C-termini of the peptide are charged (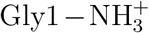 and Gly18 −COO^−^, respectively). To mimic experimental conditions, 100 mM of NaCl are added to the system. The ELPs are solvated in a cubic box with the initial sizes *l*_box_ = 6 nm. This box size ensures that the chains do not interact with their periodic images. All simulations are conducted using the software package GROMACS 2020.2.^42,43^ Based on former work on the peptide sequence GVG(VPGVG)_*n*_,^13,44^ we choose the following temperatures to investigate the LCST behavior: *T* = 288 K, 298 K, 318 K, 338 K, and 350 K, where the inverse transition temperature *T*_t_ is expected to be around room temperature for the sequence of the similar length and composition. ^31,45^ Each simulation is composed of the following steps. First, the entire system is subjected to energy minimization using the steepest descent method with the maximum force tolerance level set as 500 kJ mol^−1^ nm^−1^. Next, a short equilibration step in the *NpT* ensemble at the desired temperature *T* and at a pressure of 1 bar is performed for 200 ps. The positions of the peptide’s atoms are restrained during this step. Finally, a main run in the *NpT* ensemble is performed for 2000 ns for single-chain and multi-peptide simulations at temperatures *T >* 288 K. At a temperature of *T* = 288 K we increase the simulation time up to 4000 ns. These simulation times exceed the relaxation time of the peptide by two orders of magnitude, confirming sufficient sampling time. We use the time step of 2 fs since all bonds are constrained via the LINCS method. ^46^ Pressure coupling is maintained through a Parrinello-Rahman barostat^47^ with a time constant of 2.0 ps, while temperature coupling is achieved via a stochastic velocity rescaling thermostat with a time constant of 1.0 ps.^48^ Using the particle mesh Ewald method,^49^ long-range electrostatic interactions are computed with a cut-off of 1.2 nm. The van der Waals cut-off is also set to 1.2 nm.

We also investigate multi-chain systems employing the same simulation protocol. We perform simulations of *n*_ch_ = 2, 4, 6, and 10 peptide chains at three temperatures (*T* = 288 K, 298 K, and 350 K). The resulting peptide concentration in a simulation box varies from *c*_pept_ = 23 mg/ml (*n*_ch_ = 2) to *c*_pept_ = 115 mg/ml (*n*_ch_ = 10). The concentrations studied are far below the so-called overlap concentration ^50^ 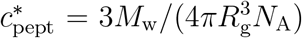 at which separatedchains start to overlap. For the ELP sequence studied here 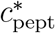 is approximately 580 mg/ml.

Additionally, we examine the interactions between two ELPs at five temperatures *T* = 288, 298, 318, 338 and 350 K employing umbrella sampling (US) ^51–53^ in which a series of configurations are generated along a reaction coordinate between two interacting species. For our system, we choose the distance between the centers of mass (CM) of the ELPs, Δ*r*, as a reaction coordinate, which is fixed over a simulation run, e.g. a sampling window, by a harmonic potential. We probe the distances between two peptides up to Δ*r* = 5.0 nm using an asymmetric distribution of the sampling windows. The window spacing *δ* is 0.1 nm for Δ*r* up to 2 nm, and *δ* = 0.2 nm for Δ*r* from 2 to 5 nm. We also vary the force constant *k* used in the umbrella potential to ensure sufficient overlap between the windows. For Δ*r <* 2 nm *k* = 1000 kJ/mol, while for larger distances *k* = 400 kJ/mol. Additionally, at temperature *T* = 288 K we increase *k* up to 1500 kJ/mol for Δ*r <* 0.5 nm to achieve an adequate sampling. We conduct three independent US calculations, where each window is sampled over a 20 ns MD run resulting in a total of 2100 ns of the simulation time for each *T*. To obtain a final curve of the potential of mean force (PMF) from our computations we use the weighted histogram analysis method.^54^ We also add the entropic correction 2*k*_B_*T* log Δ*r* and shift the curves so that the potential goes to zero at large distances.

To compute dynamical properties of the systems such as the mean-squared displacement (MSD), we conduct simulations in the *NVT* ensemble at the averaged density ⟨*ρ*(*T*)⟩ with the simulation length set to 20000 ns. This choice of the ensemble is motivated by (i) avoiding artificial displacement present in the *NpT* ensemble - position rescaling required for keeping particles in a box the size of which changes during a run; (ii) the recent finding that a heuristic unwrapping scheme employed in simulation packages including GROMACS results in unphysical trajectories due to the incorrect consideration of the fluctuating values of the box size.^55^

We analyze trajectories using in-house scripts as well as GROMACS analysis tools such as the implementation of the DSSP algorithm^56,57^ for evaluating peptide secondary structure, the number of hydrogen bonds, the MSD, a python library MDTraj^58^ for computing pairwise distances, the radius of gyration, contact maps. We used PyEmma software^59^ for estimating Markov State Models for a peptide chain.

## Results and Discussion

### LCST-like behavior at the single chain level

First, we investigate the conformational properties of a single peptide across the temperature range studied. We compute the radius of gyration *R*_g_ of the chain as

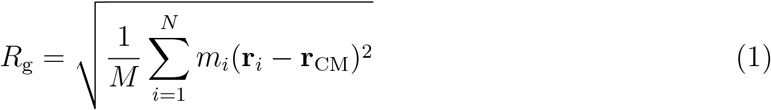

where 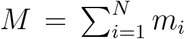 is the total mass of the peptide consisting of *N* atoms, and 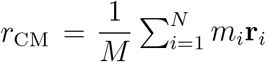 the position of the center of mass (CM) of the chain. In Figure 1(a), we present the probability density function of the radius of gyration. For *T <* 300 K we report the broad size distribution of chain sizes (large fluctuation in the size of the chain). As temperature increases, a shift in distribution towards more compact conformations is observed indicating a possible hydrophobic collapse of the chain. However, as the chain length is only eighteen amino acids, this shift in *R*_g_ is moderate.

**Figure 1:**
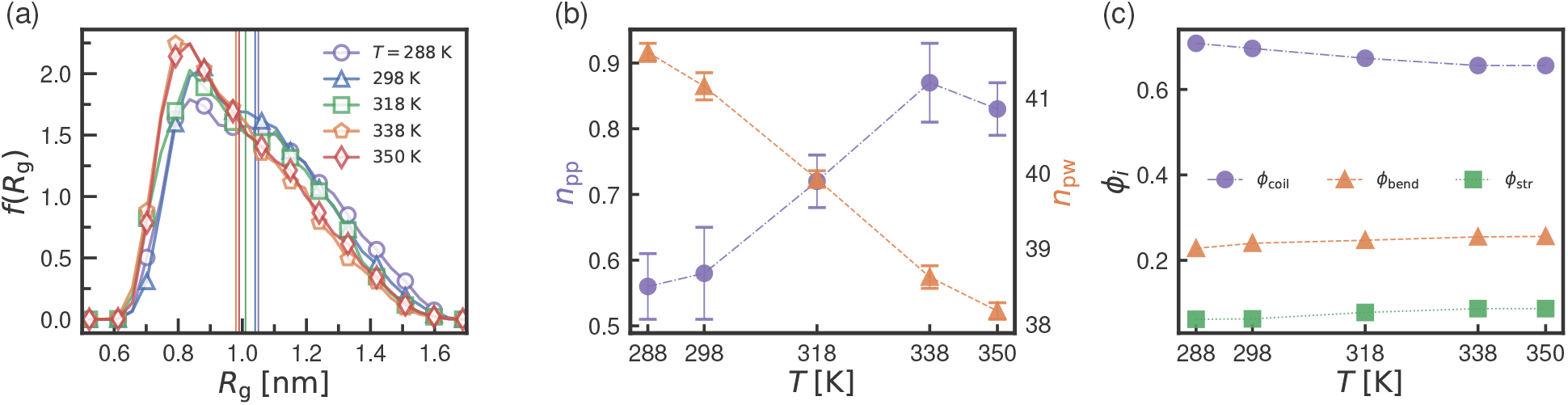
Structural properties of a single GVG(VPGVG)_3_ peptide as a function of temperature. (a) Probability density function of the radius of gyration *f* (*R*_g_) for a single ELP at five *T* investigated, vertical lines correspond to ⟨*R*_g_(*T*)⟩; (b) number of intrapeptide hydrogen bonds *n*_pp_ (left purple axis) and number of the hydrogen bonds between peptide and water molecules *n*_pw_ (right orange axis) as a function of *T*; (c) fraction of residues attributed to the following secondary structure motifs: coil *ϕ*_coil_, bend *ϕ*_bend_, and structure *ϕ*_str_ (turn+helix+*β*-bridge/sheet) as function of *T*. Dashed and dash-dotted lines are a guide to the eye.

The LCST behavior of ELPs is strongly linked to peptide-water and peptide-peptide interactions. In Fig. 1(b) we plot the number of intrapeptide hydrogen bonds *n*_pp_ and the number of hydrogen bonds between the peptide and water molecules *n*_pw_. There is a clear increase in *n*_pp_ with *T*, whereas the number of hydrogen bonds between water and the peptide *n*_pw_ decreases, indicating a change of interactions in the system and a hydrophobic collapse of the chain. A similar change in the hydrogen bond network was reported in a computational study by Rousseau et al. ^60^ The secondary structure of the peptide is mostly composed of irregular elements with a fraction of approximately 0.90 including coils and bends. Ordered structures, predominantly turns, make up the remaining. The composition of the secondary structure changes only slightly as temperature increases, where the fraction of bends and structured elements, i.e. predominantly turns, increases by a few percent at the expense of residues belonging to coil-motifs at lower *T* shown (see Fig. 1(c)). Our findings are in line with former works on ELPs and IDPs, where the hydrophobic collapse was linked to the formation of the secondary structures such as helix and turns. ^61–63^ To conclude, at the level of a single ELP we observe a moderate shift towards a more compact chain conformation as temperature increases indicating LCST-like behavior.

### Markov state model for a single peptide

As seen in Fig. 1(a), the distribution of the chain size resembles a bimodal distribution at *T* = 288 K. We attempt to characterize the metastable states visited by the peptide, their population and the transition rates between them by employing a Markov State Model (MSM) analysis.^64,65^ First, we select the backbone torsion angles as the set of variables (e.g. input features) to describe the conformational space. All configurations in a trajectory are aligned to the first frame to remove the influence of translational and rotational motions of the peptide. To decrease the dimensionality of the input data we use time-lagged independent component analysis (TICA)^66^ to find a low-dimensional representation reflecting the slowest conformational motions of the chain. After transforming the data into the representation with reduced dimension (i.e. TICA representation), the conformational space is discretized into microstates using the *k*-means clustering method, ^64^ resulting in 30 microstates. To select the optimal lag time *τ* for the model construction, MSMs are generated for several values of *τ* and the dependence of their implied timescales (ITs) *t*_*i*_ on *τ* is presented in Fig. 2(a), where *t*_*i*_ approximates the decorrelation time of the *i*^*th*^ process and is computed using the eigenvalues *λ*_*i*_ of the MSM transition matrix as *t*_*i*_(*τ*) = −*τ/* log *λ*_*i*_(*τ*). When ITs become independent on the MSM lag time (*τ*), we conclude that ITs have converged and select the smallest value of *τ* to maximize MSM resolution.^67^

**Figure 2:**
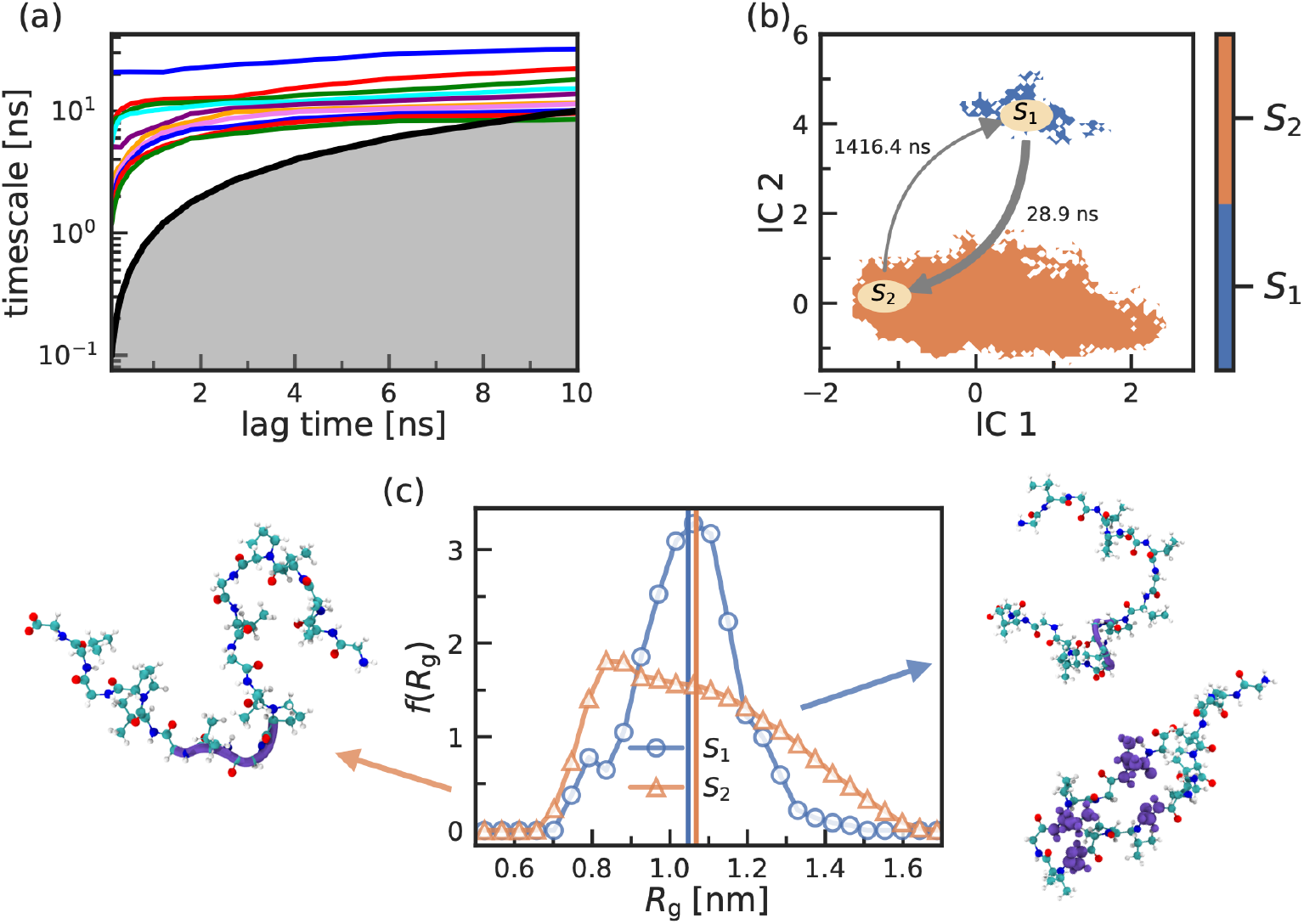
MSM analysis of a single peptide trajectory(*t*_sim_ = 4*µs*) at *T* = 288 K. (a) Implied timescales (ITs) as a function of the MSM lag time, where the black solid line corresponds to equality between ITs and the MSM lag time; (b) projection of the coarse-grained MSM (*τ* =5 ns, two macrostates) to IC1 and IC2, the mean first passage times between the states are shown as well; (c) probability density function of the radius of gyration *R*_g_ two MSM states determined, vertical lines correspond to ⟨*R*_g_⟩. Representative snapshots per state are shown as well.

In our system, ITs level off at a lag time of around 5 ns. We note that there is a distinct separation (gap) between the slowest (blue curve) and the remaining ITs (other colored curves), indicating apparent two-state dynamics. Previously, a two-state description of conformational changes was also adopted in the context of a shorter ELP sequence. ^60^ Next, we classify the 30 microstates into two macrostates with a lag time *τ* = 5 ns using the Perron Cluster Analysis method. ^68^ We validate that the final MSM satisfies the Markovian assumption by performing a Chapman-Kolmogorov (C-K) test. The C-K test is calculated using the equation *T*(*kτ*) ≈ *T*^*k*^(*τ*), where *T*(*τ*) is the transition probability matrix with a selected lag time and *k* is an integer number of steps. (see Fig S1.) In Fig. 2(b) we show the projection of the resulting MSM on the first two independent components IC1 and IC2. The trajectory mostly populates the second state *S*_2_ which contains 98 % of the trajectory, while *S*_1_ is clearly a short-lived state. To access the conformational properties of the peptide in each state, we first compute the probability density function of the radius of gyration *f* (*R*_g_) shown in Fig. 2(c). In the case of *S*_1_, *f* (*R*_g_) is characterized by a single peak with the mean value ⟨*R*_g_⟩ = 1.05 nm. The distribution for *S*_2_, however, is rather broad and resembles one from Fig.1(a). This is expected since *S*_2_ makes up almost the full trajectory. As evident from *f* (*R*_g_) for state *S*_2_, we suspect that it possibly contains several states which cannot be separated due to the strong overlap of the associated timescales (see Fig. 2 (a)), indicating the highly dynamical nature of ELPs. Next, we compute the secondary structure for each state. At first glance, the composition of the secondary structure is similar. For both states, the probability that an amino acid belongs to irregular elements (i.e. coils) is the highest (approx. 70 %), followed by bend structures (approx. 20%) and the hydrogen-bonded turn structures (approx. 5 %) shown in Fig. S2. The striking difference between the two states steamed from the formation of 3-helix and isolated *β*-bridges seen for *S*_1_. When averaging over all (eighteen) residues the probabilities of these structures might seem negligible. However, our calculations revealed that about 30% of configurations in *S*_1_ contain a 3-helix structure formed by Gly11-Val12-Gly13 residues. Mostly Val-Gly residues that are separated along the chain but spatially close form *β*-bridges. Such conformations make up 18 % of *S*_1_. The simultaneous formation of *β*-bridges and 3-helix structures does not occur, as the same or neighboring amino acids participate in structure formation. That is in contrast to *S*_2_ where only 0.4 % of the conformations show the formation of 3-helix mostly by Gly6-Val7-Gly8 and Gly11-Val12-Gly13 triplets, and *β*-bridges making 7 %. Thus, *S*_1_ is a state rich in the secondary structure with a narrow distribution of the radius of gyrations, while *S*_2_ might be described as a disordered state.

### Multi-chain systems

#### Potential of mean force

We begin our discussion of multi-peptide systems by investigating the interactions between two ELPs at five temperatures *T* = 288 K, 298 K, 318 K, 338 K and 350 K. In Figure 3, we show the potential of mean force (PMF), *w*(Δ*r*), as a function of the separation distance between CMs of the ELPs. For distances Δ*r >* 3.0 nm the interaction between two peptides goes to zero, thus we only plot the curves until Δ*r* = 3.2 nm. It is clear that at low temperature (*T* = 288 K) peptides experience repulsive interaction at short distances, while at higher temperatures (*T* = 350 K), we report a weak attraction between the chains. This confirms the LCST-like behavior of our system. To our knowledge, this is the first time that the PMF is calculated for evaluating interactions between two ELPs.

**Figure 3:**
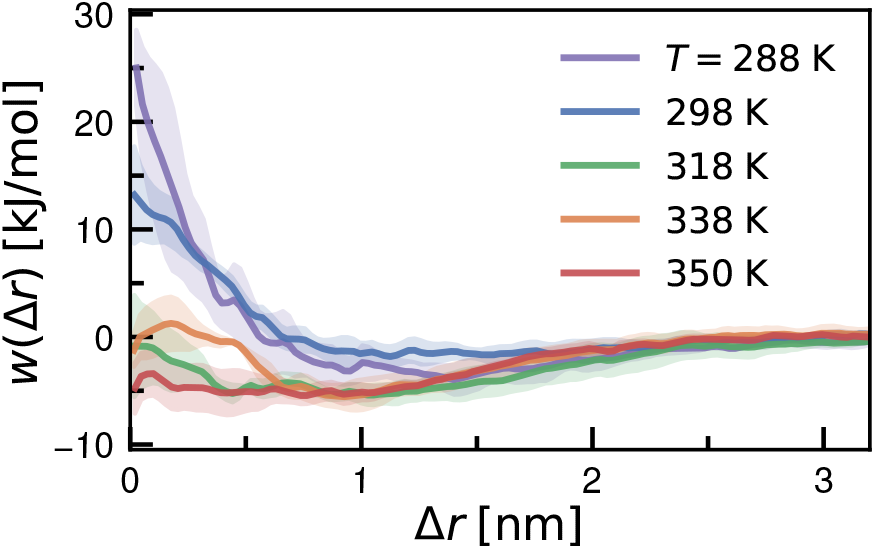
Potential of mean force *w*(Δ*r*) for a pair of ELP chains as a function of temperature. Shaded areas represent the error bars, calculated as the standard error of the mean from three independent realizations.

#### Contact formation in multi-chain systems

Furthermore, we investigate systems composed of two, four, six, and ten peptide chains at three temperatures *T* = 288 K, 298 K, and 350 K which represent temperatures below, around, and above LCST, respectively. The resulting peptide concentrations correspond to dilute solutions. (see sec. Simulation Model and Methods)

We investigate interpeptide interactions by first computing contact maps between the chains. We define two peptides as being in contact if the minimal distance between any two atoms from any two residues falls below the cutoff radius of 0.5 nm. The contact probability is normalized by the number of possible peptide pairs and the number of snapshots in which at least one aggregate is formed. In Fig. 4 (a)-(c) we show contact maps for the system at *c*_pept_ = 46 mg/ml (*n*_ch_ = 4). Similar results are obtained for other concentrations. We find that a dimer could have both an anti-parallel and a parallel orientation as shown in Fig. 4 (a) and (b), respectively. At higher *T*, more amino acids participate in contacts while the probability of contact formation slightly increases as seen in the upper panel of Fig. 4. Based on all *c*_pept_ investigated, we conclude that the most commonly found amino acid pairs which form interpeptide contacts are: Val12-Val12, Val7-Val7, Val9-Val9, and Val7-Pro10. Similarly, the amino acids that are most likely to participate in contact formation are Val12, Val7, Pro10, Val9, and Val 14. Thus, valine amino acids located in the middle of the chain drive inter-peptide contact formation.

**Figure 4:**
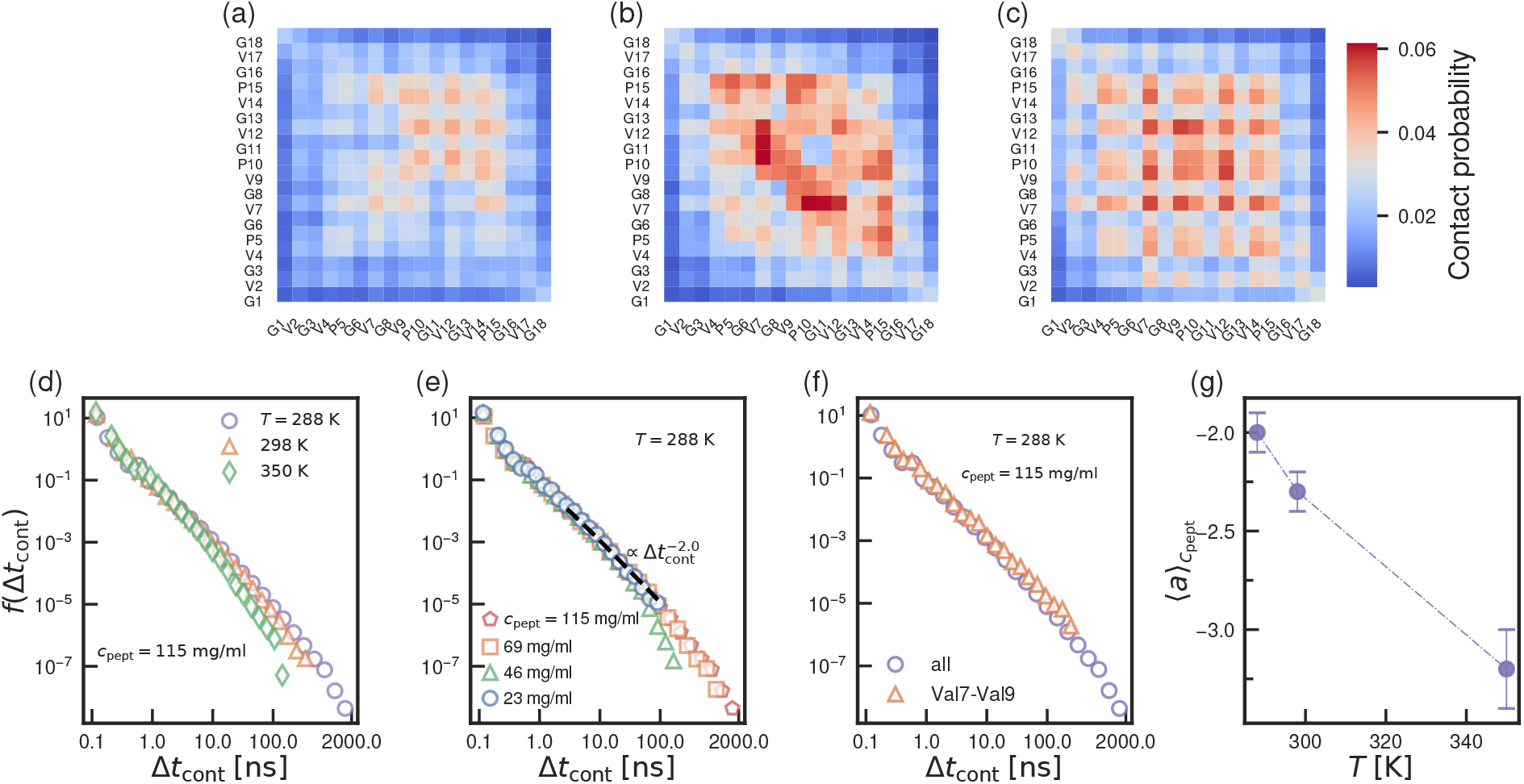
Contact formation in multi-peptide systems. (a), (b), and (c) Interpeptide contact maps for the system at *c*_pept_ = 46 mg/ml at temperatures *T* =288 K, 298 K, and 350 K, respectively. A cutoff of 0.5 nm is used to define a contact between atoms. Contact maps are constructed using the same scale; (d) and (e) probability density function of contact time *f* (Δ*t*_cont_) as a function of *T* for *c*_pept_ = 115 mg/ml and as a function of *c*_pept_ for *T* = 288 K, respectively; (f) probability density function of contact time *f* (Δ*t*_cont_) for all contacts *vs* the most likely contact forming pair (Val7-Val9) for the system at *c*_pept_ = 115 mg/ml and *T* = 288 K; (g) fitting exponent ⟨*a*⟩_*c*pept_ to *f* (Δ*t*_cont_) as a function of *T*.

To probe the utility and generality of our findings, we generate six new peptide sequences where we modify Val at the position of the guest residue at positions 7, 12, and 17 with more hydrophilic amino acid alanine (Ala) and Gly which is a residue that suppresses aggregation in IDPs.^69^ Experimentally, it was also shown that using both sequence modifications results in the shift of the LCST temperature to values above 60 ^◦^ C.^70^ We investigate new sequences at two temperatures *T* =288 K, and 350 K, i. e. below and above LCST for the original sequence, and for three concentrations investigate single-chain (*c*_pept_ = 11.5 mg/ml), two-chain (*c*_pept_ = 23 mg/ml), and ten-chain systems (*c*_pept_ = 115 mg/ml). At a single-chain level, the modification of the sequence shows only a minor influence on the chain dimensions (shown in Fig. S3). The strongest change we observe for the sequences where Ala was introduced at the 12 and 17 positions. Next, we compare the contact formation in these systems with the original sequence. We find that for both peptide concentrations and at both temperatures, the probability of forming contacts is reduced for all new sequences as shown in Fig. 5 and Fig. S4 for Ala and Gly modifications, respectively. The sequence modification in the 7 or 12 positions with Ala or Gly shows the strongest impact on the formation for both concentrations, where the value of the contact probability drops by up to five times. The effect on contact formation by sequence modification in the 17 position is milder, as this residue does not actively participate in contacts. Thus, we highlight the important role of Val residues in the middle of the chain in contact formation.

**Figure 5:**
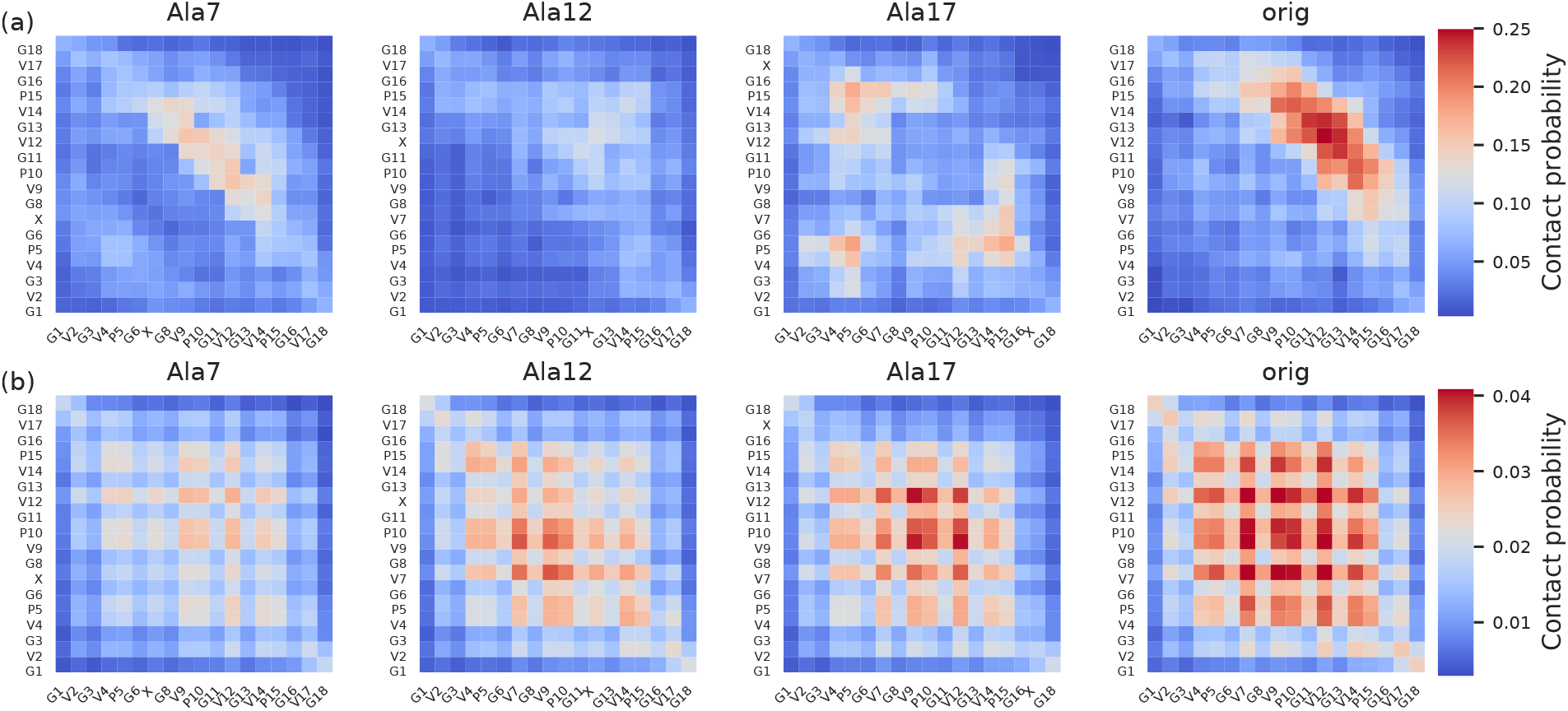
Contact formation in sequences modified with alanine. (a) and (b) Interpeptide contact maps for the system at *c*_pept_ = 23 mg/ml at temperatures *T* =288 K and *c*_pept_ = 115 mg/ml and *T* =350 K, respectively. The sequence modification is indicated by a label on top of each contact map. Label “orig” corresponds to the originally studied sequence. A cutoff of 0.5 nm is used to define contact between atoms. Contact maps are constructed using the same scale.

Additionally, we analyze the time scales of contact events and their dependence on temperature and peptide concentration. In Fig. 4 (d) we show the probability density of contact lifetime *f* (Δ*t*_cont_) as a function of *T* at a concentration *c*_pept_ = 115 mg/ml (*n*_ch_ = 10). Most of the contacts are short-lived highlighting the highly transient nature of dimers, i.e. for Δ*t*_cont_ ≤ 2 ns their statistics are independent of *T*, while for longer Δ*t*_cont_ the curves deviate from each other and can be fitted by a power law decay with a different exponent. For lower *T*, we detect Δ*t*_cont_ up to 2 *µ*s, while at higher *T* values of Δ*t*_cont_ are limited by hundreds of nanoseconds, indicating increased mobility of amino acids. Additionally, *f* (Δ*t*_cont_) decays faster with temperature. The peptide concentration does not affect the slope of *f* (Δ*t*_cont_) as shown in Fig. 4 (e), but due to increased sampling at higher concentrations, longer Δ*t*_cont_ can be observed. For Δ*t*_cont_ in the range of a few to hundred nanoseconds, we find a power law

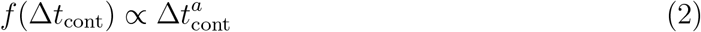

with an exponent *a* that shows no significant dependence on ELP concentration is approx-imately -2 at *T* = 288 K and decreases to below -3 for *T* = 350 K. Thus, we find that temperature strongly affects the distribution of contact lifetimes since the exponent averaged over peptide concentrations ⟨*a*⟩_*c*pept_ decreases with *T* as presented in Fig. 4 (g). Similar observations are valid for the modified Ala and Gly sequences and are presented in Fig. S5. The individual contacts at higher temperatures thus become more transient, even though the total number of contacts increases (see Fig. S6), suggesting a cooperative nature of aggregation.

As we identified amino acid pairs that most often participate in contact events, we compare the distribution *f* (Δ*t*_cont_) for such a pair, i.e. Val7-Val9, against the one that includes all contact lifetimes for a system *c*_pept_ = 115 mg/ml at *T* = 288K. The corresponding curves overlap as seen in Fig. 4, (f) indicating that there are no significant differences between the contact time distributions.

To rationalize the observed power law dependence of the contact time Δ*t*_cont_ on *T* shown in Fig. 4 (e), we use the well-known “trap mode” ^71^ that allows one to obtain a nontrivial power law dependency based on two very simple ingredients: (i) the Arrhenius equation provides the relation between the activation energy *E* and a timescale *τ* of an activated process *τ* = *τ*_0_ exp(*E/k*_B_*T*), where *τ*_0_ is a characteristic time and *k*_B_ is the Boltzmann constant; (ii) *E* follows an exponential distribution near the value *E*_0_ as 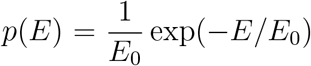. Usinga change of variable for the distribution of the lifetime *f*(*τ*): *f*(*τ*)*dτ* = *p*(*E*)*dE*, one easily obtains

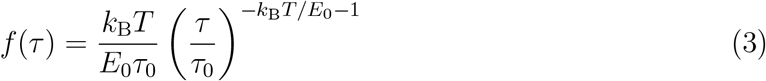

where the exponent *a*_th_ = −*k*_B_*T/E*_0_ − 1. Thus, *a*_th_ is inversely proportional to *T* which is consistent with our findings. However, we were unable to fit 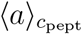(*T*) assuming that *E*_0_ is a constant. We find instead that the ratio *E*_0_*/k*_B_*T* decreases from 1.0 to 0.6 where *T* = 288 K is used. Thus, the interaction energy between two peptides becomes less repulsive with temperature, which is in accordance with the LCST-like behaviour of ELPs. Similar to our findings, M. Rickard et al^72^ also reported a power law decay of protein-protein contact lifetimes in a crowded environment.

#### Structural properties of the peptides and aggregates

Next, we investigate the structural properties of multi-chain systems. As mentioned above, we use the cutoff distance (0.5 nm) to define an aggregate. The radius of gyration computed for chains that belong to an aggregate is shown in Fig. 6 (a). For comparison, we also add *f* (*R*_g_) for the single-chain system. We observe a broad distribution *f* (*R*_g_) for systems with several chains, indicating that a peptide can assume both compact and more stretched conformations when being part of a cluster. Our findings are in line with a numerical study by S. Rauscher et al,^34^ where the expansion of ELPs was reported when the peptides formed aggregates.

**Figure 6:**
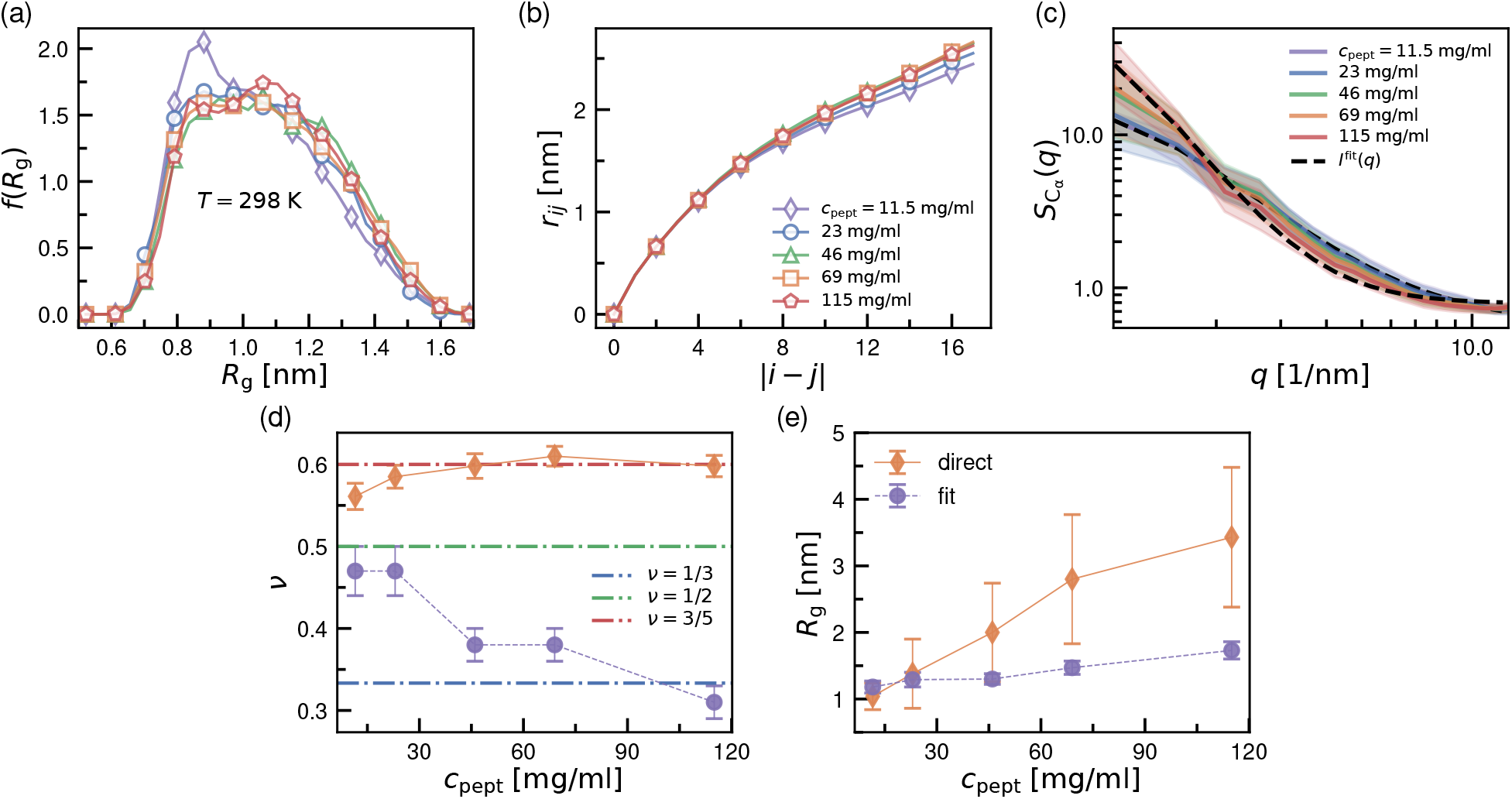
Structural properties of ELPs as a function of the concentration at *T* =298 K. (a) Probability density function of the radius of gyration *f* (*R*_g_) of a chain partitioning into an aggregate; (b)root-mean-square internal distance *r*_*ij*_ between *C*_*α*_ separated by |*i*− *j*| amino acids along the peptide chain; (c) static structure factor 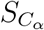 (*q*), black dashed lines represent fits to the curves using the scattering function of a single chain *I*(*q*), while shaded areas represent the error bars, calculated as the standard error of the mean value across a trajectory; (d) Flory exponent *ν* as a function of *c*_pept_ obtained from the fits of the internal distances *r*_*ij*_ (orange diamonds, direct) and fits to the static structure factor 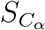 (*q*) with *I*(*q*) expression (Eq. 5, purple circles, fit); (e) radius of gyration *R*_g_ as a function of *c*_pept_ computed for the largest aggregates (orange diamonds) and obtained from the fits to the static structure factor 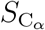 (*q*) with *I*(*q*) expression (Eq. 5, purple circles). Note that (a) and (b), (d) and (e) share the same legend.

As another structural characteristic, we compute the root-mean-square internal distance *r*_*ij*_ between *C*_*α*_ atoms separated by |*i* − *j*| amino acids along the peptide chain, only considering chains that form aggregates. As shown in Fig. 6 (b), *r*_*ij*_ slightly increases with peptide concentration, evidencing chain expansion. The same outcome holds for all *T* investigated here. We only present the value at *T* = 298 K for the sake of brevity.

As a separate measure for structural properties of the entire system, we analyze the structure factor 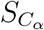 (*q*) considering positions of *C*_*α*_ atoms only shown in Fig. 6 (c):

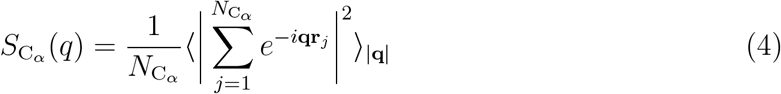

where **q** is the scattering wavevector of a length |**q**| = *q*, **r**_*j*_ is position of *C*_*α*_ atom, and 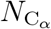 is the total number of *C*_*α*_ atoms in the system. We fit 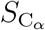 (*q*) curves with the analytic scattering function for a single Flory chain commonly employed in experiments ^44,73^

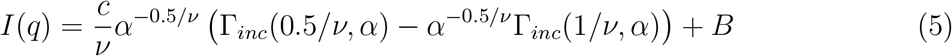

where *c* is a constant that depends on peptide concentration, 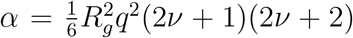, Γ(*x*) is the gamma function, and *B* is the background. In Fig. 6 (d) we summarize the Flory exponent *ν* obtained from fitting *r*_*ij*_ = *a*|*i* − *j*|^*ν*^ (direct) as well as derived from Eq. 5 (fit) at *T* = 298 K. At all peptide concentrations, *ν* values extracted from the fits to 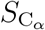 (*q*) strongly underestimate the chain dimensions, predicting its conformation to be between an ideal (*ν* = 1*/*2) and a collapsed (*ν* = 1*/*3) chain. In contrast, real *ν* values calculated from internal distances describe chain conformation as an ideal or a swollen (*ν* = 3*/*5) coil.

We also compare the radius of gyration *R*_g_ computed for the largest clusters formed (=*n*_ch_) with the values extracted from the fits (see Fig.6 (e)). As for the Flory exponent, directly computed *R*_g_ values exceed the ones extracted from the fits for systems containing more than two chains. We conclude that both methods yield similar values for *R*_g_ and *ν* only for single- or two-chain systems (*c*_pept_ = 11.5 and 23 mg/ml, respectively). This finding is not surprising, since Eq.5 is derived for a single chain. Even though the peptide concentrations are way below the overlap concentration, and the concentration range corresponds to dilute conditions, the entangled system does not represent a single linear chain, and caution is needed when utilizing functions derived from single-chain systems to interpret results. While the values for *R*_*g*_ are completely erroneous, the Flory exponent can be understood approximately as follows: when mapping a system of interacting chains (i.e. forming aggregates) during the fit onto an effective chain, this effective chain obviously has a more compact structure than the individual extended chains. The increasing deviation between the correct single-chain exponent and the effective Flory exponent for the full system (Fig. 6(d)) suggests a significantly increased effect of chain clustering with increasing concentration.

#### Dynamical properties of peptides

We now turn to the discussion of the dynamical properties of the ELPs. We remind the reader that our simulations are conducted in the *NVT* ensemble since it is more suitable for calculations (see sec. Simulation Model and Methods). We first compute the mean squared displacement (MSD) ⟨Δ*r*^2^⟩ of individual peptides using the positions of the central *C*_*α*_ ^34^ across *c*_pept_ at *T* = 298 K shown in Fig. 7 (a). As one could expect, ELPs become less mobile as *c*_pept_ in the system increases for all *T* investigated. We extract the diffusion coefficient of individual chains *D* by fitting the MSD with the expression ⟨Δ*r*^2^⟩ = 6*Dt*. Additionally, we compute the diffusion coefficient in the dilute limit using the Stokes-Einstein equation *D*_0_(*T*) = *k*_B_*T/*[6*πη*(*T*)*R*_h_(*T*)], where *η*(*T*) is the viscosity of the water model,^36^ 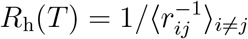 is the hydrodynamic radius of the peptide estimated through the Kirkwood-Riseman equation using internal distances *r*_*ij*_ from single-peptide runs.^74^ In Fig. 7 (b) we present the ratio *D*_0_*/D* as a function of the peptide volume fraction 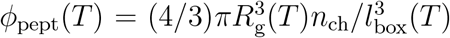, where we use values of *R*_g_(*T*) and *l*_box_(*T*) from single-peptide simulations. With increasing *ϕ*_pept_ and in particular for *ϕ*_pept_ ≥ 0.1, the diffusion is slowed down, as intuitively expected for a more packed system. Importantly, increasing the temperature leads to a slowdown of the diffusion, which is in line with the increased attractive interaction between the peptide chains. However, it has to be stressed that this temperature-induced slowdown of a factor smaller than two at a fixed volume fraction (*ϕ*_pept_ ≤ 0.1) is rather mild, which reflects the transient nature of interpeptide contacts.

**Figure 7:**
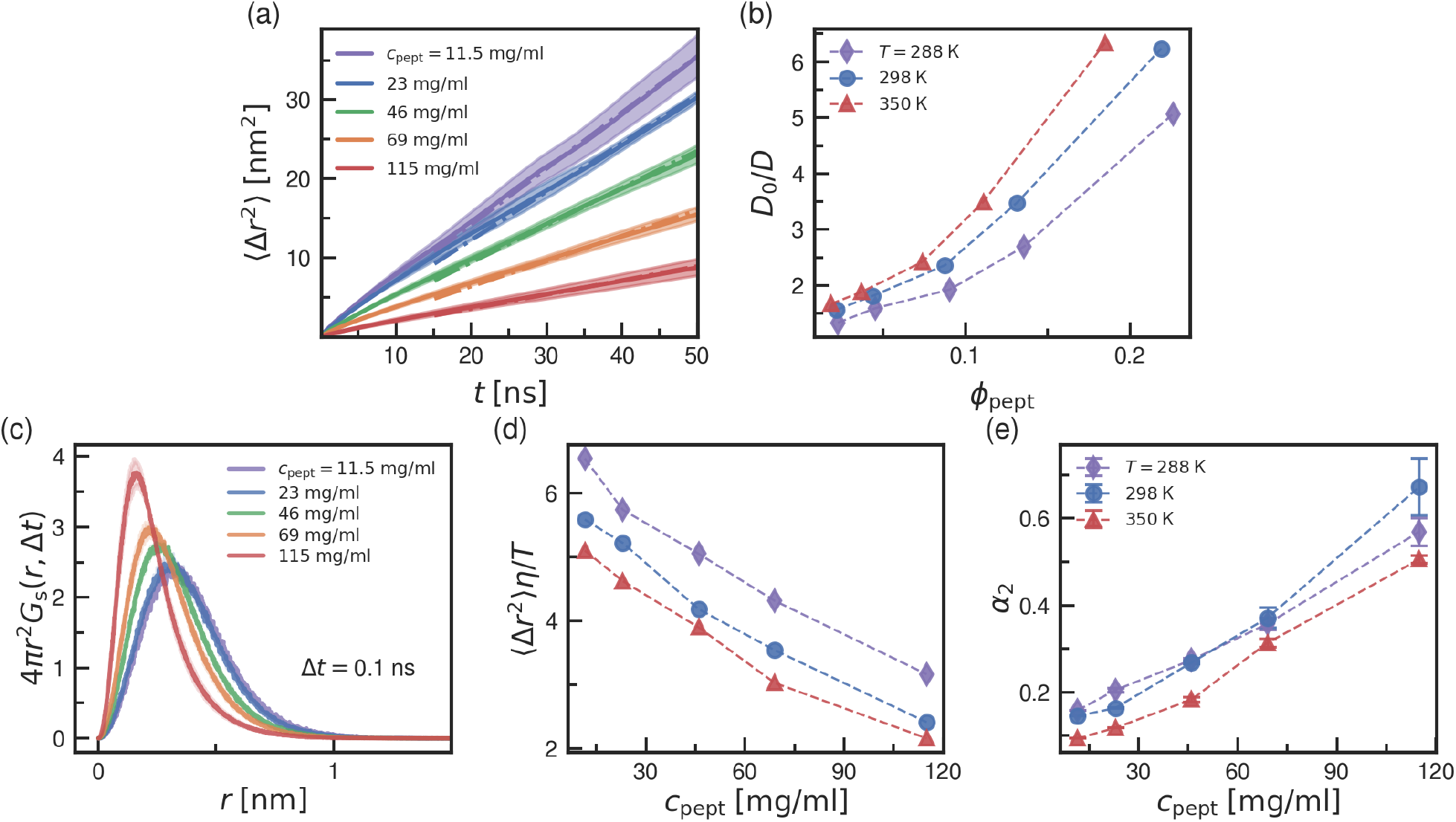
Dynamical properties of ELPs. (a) Mean squared displacement (MSD) ⟨Δ*r*^2^⟩ of the central *C*_*α*_ atom at *T* = 298 K. Shaded areas represent the error bars, calculated as the average MSD for multi-chain simulations and the mean error between diagonal components for the single peptide simulations. Dash-dotted lines represent the fits to the curves using the expression ⟨Δ*r*^2^⟩ = 6*Dt*; (b) ratio of the diffusion coefficients *D*_0_*/D* as a function of the volume fraction *ϕ*_pept_; (c) the van Hove self-correlation function *G*_s_(*r*, Δ*t*) computed using the backbone hydrogen atoms at *T* = 298 K for the time windows Δ*t*=0.1 ns; (d) ⟨Δ*r*^2^⟩*η/T* [10^−17^m^2^Pa · s*/*K] as a function of the peptide concentration *c*_pept_, where ⟨Δ*r*^2^⟩ is computed for the time window Δ*t*=0.1 ns; (e) non-Gaussian parameter *α*_2_ as a function of the peptide concentration *c*_pept_, *α*_2_ is also computed for Δ*t*=0.1 ns. For the graphs where the error bars are not shown, the standard error is smaller than the symbol size.

Furthermore, we calculate the van Hove correlation function that provides information about the type of dynamics in the system. In particular, the self-part of the van Hove correlation function *G*_s_(*r*, Δ*t*) describes the probability that an atom has moved from its initial position to a distance *r* after the time interval Δ*t*

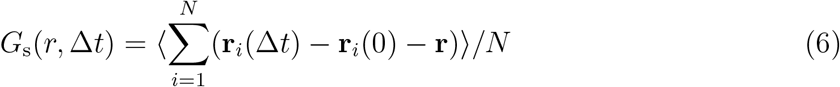

where **r**_*i*_ is the position of the *i*^*th*^ atom, **r** is a position in space with a distance |**r**| = *r* from the origin, and *N* is a total number of atoms. For an ideal fluid where particles undergo Brownian motion *G*_s_ (*r*, Δ*t*) becomes a Gaussian function: 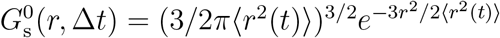. Also, *G*_s_(*r*, Δ*t*) is the Fourier transform of the intermediate scattering function *F*_*s*_(**q**, *t*) that can be obtained experimentally by incoherent quasi-elastic neutron scattering measurements.^75^ In our calculations we consider the positions of hydrogen atoms belonging to the backbone for future comparison with experimental data. In Figure 7 (c) we show *G*_s_(*r*, Δ*t*) for time intervals Δ*t* = 0.1 that reflect the internal motion of the chain at *T* = 298 K for the full concentration range. As expected, the hydrogen motion is more constrained at high *c*_pept_. To get a better view on the fast dynamics of the chain, we additionally compute MSD normalized by the temperature effects ⟨*r*^2^(*t*)⟩*η/T* for the same time window (Δ*t* = 0.1) (see Fig. 7 (d)). We observe that the internal chain motion is slowed down by the increase in peptide concentration and temperature. The latter highlights stronger interchain interactions at higher temperature that is consistent with the LCST phase-behavior. To characterize deviations of atom motion from Brownian motion, i.e. difference between *G*_s_(*r*, Δ*t*) and 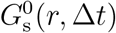, we compute the so-called “non-Gaussian” parameter *α*_2_(Δ*t*) = 3⟨*r*^4^(Δ*t*)⟩*/*5⟨*r*^2^(Δ*t*)⟩_2_ − 1 forΔ*t* = 0.1. By construction, *α*_2_(Δ*t*) = 0 for a Brownian process. As seen in Fig. 7 (e), atom displacements deviate from the Gaussian limit as peptide concentration increases for all *T* investigated.

## Conclusions

We investigated in-depth structural and dynamical properties of aqueous solutions of the ELP GVG(VPGVG)_3_ as a function of the peptide concentration in a temperature range corresponding to their LCST-like phase behavior. We used a state-of-the-art force field (i.e. the AMBER99SB-ILDN force field for the peptide and the TIP4P-D water model) that is suitable for modeling IDPs. On a single-chain level, there is a signature of a hydrophobic collapse already as temperature increases. However, due to the short length of the sequence investigated, the chain size decreases only slightly. Using Markov State Modeling analysis, we characterized the conformational ensemble explored by the peptide using a two-state representation, in which one state is rich in secondary structure, while the second one could be described as disordered. For two-peptide systems, the potential of mean force between two chains across a temperature range shows a change in interaction consistent with LCST-behavior. In multi-chain systems, peptides within aggregates are more extended compared to a single chain in solution. Contacts formed between peptides are highly dynamical and become more transient as the temperature is raised. Contact lifetime distributions are well described by a power law decay. Central Val residues drive contact formation, especially at lower temperatures. Replacement of these residues with more hydrophilic residues Ala or weaker interacting Gly strongly suppresses contact formation. Peptide dimensions in a single and multi-chain systems lay between an ideal and a swollen coil conformation when using real space coordinates. Surprisingly, extracting the Flory exponent from the static structure factor suggests a globular conformation. The two methods approaches are in agreement only when applied to extremely dilute conditions 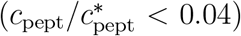 and disagree for high concentrations as the interchain interactions become important. Both the long-time diffusion and the internal chain motion are strongly slowed down by increasing the concentration while an increase in temperature only contributes a mild additional slowdown. We believe these results can be valuable for designing any application which employs thermo-responsive polymeric materials.

## Associated content

### Supporting information

Chapman-Kolmogorov test for the MSM constructed for a GVG(VPGVG)_3_ peptide; the secondary structure of a peptide resolved for each MSM state; distributions of the chain size for sequence-modified peptides; contact maps for glycine-modified sequences; contact lifetime distributions in modified sequences; the total number of contacts in multi-peptide systems.

## Acknowledgments

This work was performed using HPC resources (GPU-accelerated partitions of the Jean Zay supercomputer) from GENCI–IDRIS (Grant 2021 - A0100712464). JLB and NAG acknowledge the financial support of the International Research Project (IRP) “Statistical Physics of Materials” funded by CNRS. NAG also thanks PICT 2019-02257.

## References

(1) Mithieux, S. M.; Weiss, A. S. Elastin. Adv. Prot. Chem. 2005, 70, 437–461, DOI: https://doi.org/10.1016/S0065-3233(05)70013-9.

(2) Muiznieks, L. D.; Weiss, A. S.; Keeley, F. W. Structural disorder and dynamics of elastin. Biochem. Cell Biol. 2010, 88, 239–250.

(3) Ozsvar, J.; Yang, C.; Cain, S. A.; Baldock, C.; Tarakanova, A.; Weiss, A. S. Tropoelastin and elastin assembly. Front. Bioeng. Biotechnol. 2021, 9, 643110.

(4) Urry, D. W.; Urry, K. D.; Szaflarski, W.; Nowicki, M. Elastic-contractile model proteins: Physical chemistry, protein function and drug design and delivery. Adv. Drug Del. Rev. 2010, 62, 1404 – 1455, DOI: https://doi.org/10.1016/j.addr.2010.07.001.

(5) Shah, M.; Hsueh, P.-Y.; Sun, G.; Chang, H. Y.; Janib, S. M.; MacKay, J. A. Biodegradation of elastin-like polypeptide nanoparticles. Protein Sci. 2012, 21, 743–750, DOI: 10.1002/pro.2063.

(6) Varanko, A. K.; Su, J. C.; Chilkoti, A. Elastin-like polypeptides for biomedical applications. Annu. Rev. Biomed. Eng. 2020, 22, 343–369.

(7) Quintanilla-Sierra, L.; Garc’ sıa-Arévalo, C.; Rodriguez-Cabello, J. Self-assembly in elastin-like recombinamers: a mechanism to mimic natural complexity. Mat. Today Bio 2019, 2, 100007, DOI: https://doi.org/10.1016/j.mtbio.2019.100007.

(8) Nettles, D. L.; Chilkoti, A.; Setton, L. A. Applications of elastin-like polypeptides in tissue engineering. Adv. Drug Del. Rev. 2010, 62, 1479 – 1485, DOI: https://doi.org/10.1016/j.addr.2010.04.002.

(9) Floss, D. M.; Schallau, K.; Rose-John, S.; Conrad, U.; Scheller, J. Elastin-like polypeptides revolutionize recombinant protein expression and their biomedical application. Trends Biotechnol. 2010, 28, 37–45, DOI: https://doi.org/10.1016/j.tibtech.2009.10.004.

(10) MacEwan, S. R.; Chilkoti, A. Elastin-like polypeptides: Biomedical applications of tunable biopolymers. Peptide Sci. 2010, 94, 60–77, DOI: 10.1002/bip.21327.

(11) Ribeiro, A.; Arias, F. J.; Reguera, J.; Alonso, M.; Rodr’ sıguez-Cabello, J. C. Influence of the amino-acid sequence on the inverse temperature transition of elastin-like polymers. Biophys. J. 2009, 97, 312–320.

(12) Quiroz, F. G.; Chilkoti, A. Sequence heuristics to encode phase behaviour in intrinsically disordered protein polymers. Nat. Mater. 2015, 14, 1164–1171.

(13) Zhao, B.; Li, N. K.; Yingling, Y. G.; Hall, C. K. LCST behavior is manifested in a single molecule: elastin-like polypeptide (VPGVG) n. Biomacromolecules 2016, 17, 111–118.

(14) MacKay, J. A.; Callahan, D. J.; FitzGerald, K. N.; Chilkoti, A. Quantitative model of the phase behavior of recombinant pH-responsive elastin-like polypeptides. Biomacromolecules 2010, 11, 2873–2879.

(15) McPherson, D. T.; Xu, J.; Urry, D. W. Product purification by reversible phase transition followingescherichia coliexpression of genes encoding up to 251 repeats of the elastomeric pentapeptide gvgvp. Protein Expr. Purif. 1996, 7, 51–57.

(16) Tamura, T.; Yamaoka, T.; Kunugi, S.; Panitch, A.; Tirrell, D. Effects of temperature and pressure on the aggregation properties of an engineered elastin model polypeptide in aqueous solution. Biomacromolecules 2000, 1, 552–555.

(17) Iakoucheva, L. M.; Brown, C. J.; Lawson, J. D.; Obradovi’ sc, Z.; Dunker, A. K. Intrinsic disorder in cell-signaling and cancer-associated proteins. J. Mol. Biol. 2002, 323, 573– 584.

(18) Roberts, S.; Dzuricky, M.; Chilkoti, A. Elastin-like polypeptides as models of intrinsically disordered proteins. FEBS letters 2015, 589, 2477–2486.

(19) Brangwynne, C.; Tompa, P.; Pappu, R. Polymer physics of intracellular phase transitions. Nat. Phys. 2015, 11, 899–904.

(20) Reichheld, S. E.; Muiznieks, L. D.; Keeley, F. W.; Sharpe, S. Direct observation of structure and dynamics during phase separation of an elastomeric protein. Proc. Natl. Acad. Sci. USA 2017, 114, E4408, DOI: 10.1073/pnas.1701877114.

(21) Urry, D. W. Physical chemistry of biological free energy transduction as demonstrated by elastic protein-based polymers. J. Phys. Chem. B 1997, 101, 11007–11028.

(22) Rodr’ sıguez-Cabello, J. C.; Alonso, M.; Pérez, T.; Herguedas, M. M. Differential scanning calorimetry study of the hydrophobic hydration of the elastin-based polypentapeptide, poly (VPGVG), from deficiency to excess of water. Biopolymers 2000, 54, 282–288.

(23) Henze, R.; Urry, D. W. Dielectric relaxation studies demonstrate a peptide librational mode in the polypentapeptide of elastin. J. Am. Chem. Soc. 1985, 107, 2991–2993.

(24) Matt, A.; Kuttich, B.; Grillo, I.; Weißheit, S.; Thiele, C. M.; Stühn, B. Temperature induced conformational changes in the elastin-like peptide GVG(VPGVG)_3_. Soft Matter 2019, 15, 4192–4199, DOI: 10.1039/C9SM00583H.

(25) Pepe, A.; Armenante, M. R.; Bochicchio, B.; Tamburro, A. M. Formation of nanostructures by self-assembly of an elastin peptide. Soft Matter 2009, 5, 104–113.

(26) Lessing, J.; Roy, S.; Reppert, M.; Baer, M.; Marx, D.; Jansen, T. L. C.; Knoester, J.; Tokmakoff, A. Identifying residual structure in intrinsically disordered systems: a 2D IR spectroscopic study of the GVGXPGVG peptide. J. Am. Chem. Soc. 2012, 134, 5032–5035.

(27) Urry, D. W. Entropic elastic processes in protein mechanisms. I. Elastic structure due to an inverse temperature transition and elasticity due to internal chain dynamics. J. Protein Chem. 1988, 7, 1–34.

(28) Hong, M.; Isailovic, D.; McMillan, R.; Conticello, V. Structure of an elastin-mimetic polypeptide by solid-state NMR chemical shift analysis. Biopolymers 2003, 70, 158– 168.

(29) Ohgo, K.; Ashida, J.; Kumashiro, K. K.; Asakura, T. Structural determination of an elastin-mimetic model peptide,(Val-Pro-Gly-Val-Gly) 6, studied by 13C CP/MAS NMR chemical shifts, two-dimensional off magic angle spinning spin-diffusion NMR, rotational echo double resonance, and statistical distribution of torsion angles from Protein Data Bank. Macromolecules 2005, 38, 6038–6047.

(30) Reiersen, H.; Clarke, A. R.; Rees, A. R. Short elastin-like peptides exhibit the same temperature-induced structural transitions as elastin polymers: implications for protein engineering. J. Mol. Biol. 1998, 283, 255–264.

(31) Tarakanova, A.; Huang, W.; Weiss, A. S.; Kaplan, D. L.; Buehler, M. J. Computational smart polymer design based on elastin protein mutability. Biomaterials 2017, 127, 49– 60.

(32) Zhao, Y.; Kremer, K. Proline isomerization regulates the phase behavior of elastin-like polypeptides in water. J. Phys. Chem. B 2021, 125, 9751–9756.

(33) Li, N. K.; Quiroz, F. G.; Hall, C. K.; Chilkoti, A.; Yingling, Y. G. Molecular description of the LCST behavior of an elastin-like polypeptide. Biomacromolecules 2014, 15, 3522–3530.

(34) Rauscher, S.; Pomes, R. The liquid structure of elastin. Elife 2017, 6, e26526.

(35) Li, N. K.; Xie, Y.; Yingling, Y. G. Insights into Structure and Aggregation Behavior of Elastin-like Polypeptide Coacervates: All-Atom Molecular Dynamics Simulations. J. Phys. Chem. B 2021, 125, 8627–8635.

(36) Morozova, T. I.; Garc’ sıa, N. A.; Barrat, J.-L. Temperature dependence of thermodynamic, dynamical, and dielectric properties of water models. J. Chem. Phys. 2022, 156, 126101, DOI: 10.1063/5.0079003.

(37) Lindorff-Larsen, K.; Piana, S.; Palmo, K.; Maragakis, P.; Klepeis, J. L.; Dror, R. O.; Shaw, D. E. Improved side-chain torsion potentials for the Amber ff99SB protein force field. Proteins: Struct. Funct. Genet. 2010, 78, 1950–1958.

(38) Piana, S.; Donchev, A. G.; Robustelli, P.; Shaw, D. E. Water dispersion interactions strongly influence simulated structural properties of disordered protein states. J. Phys. Chem. B 2015, 119, 5113–5123.

(39) Henriques, J.; Cragnell, C.; Skepö, M. Molecular dynamics simulations of intrinsically disordered proteins: force field evaluation and comparison with experiment. J. Chem. Theory Comput. 2015, 11, 3420–3431.

(40) Henriques, J.; Skepö, M. Molecular dynamics simulations of intrinsically disordered proteins: on the accuracy of the TIP4P-D water model and the representativeness of protein disorder models. J. Chem. Theory Comput. 2016, 12, 3407–3415.

(41) Hanwell, M. D.; Curtis, D. E.; Lonie, D. C.; Vandermeersch, T.; Zurek, E.; Hutchison, G. R. Avogadro: an advanced semantic chemical editor, visualization, and analysis platform. J. Cheminformatics 2012, 4, 1–17.

(42) Lindahl,; Abraham,; Hess,; van der Spoel, GROMACS 2020.2 Source code. 2020; https://doi.org/10.5281/zenodo.3773801.

(43) Lindahl,; Abraham,; Hess,; van der Spoel, GROMACS 2020.2 Manual. 2020; https://doi.org/10.5281/zenodo.3773799.

(44) Matt, A.; Kuttich, B.; Grillo, I.; Weißheit, S.; Thiele, C. M.; Stühn, B. Temperature induced conformational changes in the elastin-like peptide GVG (VPGVG) 3. Soft Matter 2019, 15, 4192–4199.

(45) Urry, D. W.; Gowda, D. C.; Parker, T. M.; Luan, C.-H.; Reid, M. C.; Harris, C. M.; Pattanaik, A.; Harris, R. D. Hydrophobicity scale for proteins based on inverse temperature transitions. Biopolymers 1992, 32, 1243–1250.

(46) Hess, B.; Bekker, H.; Berendsen, H. J.; Fraaije, J. G. LINCS: a linear constraint solver for molecular simulations. J. Comput. Chem. 1997, 18, 1463–1472.

(47) Parrinello, M.; Rahman, A. Polymorphic transitions in single crystals: A new molecular dynamics method. J. Appl. Phys. 1981, 52, 7182–7190.

(48) Bussi, G.; Donadio, D.; Parrinello, M. Canonical sampling through velocity rescaling. J. Chem. Phys. 2007, 126, 014101.

(49) Darden, T.; York, D.; Pedersen, L. Particle mesh Ewald: An N.log (N) method for Ewald sums in large systems. J. Chem. Phys. 1993, 98, 10089–10092.

(50) Doi, M.; Edwards, S. F.; Edwards, S. F. The theory of polymer dynamics; oxford university press, 1988; Vol. 73.

(51) Patey, G.; Valleau, J. The free energy of spheres with dipoles: Monte Carlo with multistage sampling. Chem. Phys. Lett. 1973, 21, 297–300.

(52) Torrie, G. M.; Valleau, J. P. Monte Carlo free energy estimates using non-Boltzmann sampling: Application to the sub-critical Lennard-Jones fluid. Chem. Phys. Lett. 1974, 28, 578–581.

(53) Torrie, G. M.; Valleau, J. P. Nonphysical sampling distributions in Monte Carlo freeenergy estimation: Umbrella sampling. J. Comput. Phys. 1977, 23, 187–199.

(54) Kumar, S.; Rosenberg, J. M.; Bouzida, D.; Swendsen, R. H.; Kollman, P. A. The weighted histogram analysis method for free-energy calculations on biomolecules. I. The method. J. Comput. Chem. 1992, 13, 1011–1021.

(55) von Bülow, S.; Bullerjahn, J. T.; Hummer, G. Systematic errors in diffusion coefficients from long-time molecular dynamics simulations at constant pressure. J. Chem. Phys. 2020, 153, 021101.

(56) Kabsch, W.; Sander, C. Dictionary of protein secondary structure: pattern recognition of hydrogen-bonded and geometrical features. Biopolymers 1983, 22, 2577–2637.

(57) Joosten, R. P.; Te Beek, T. A.; Krieger, E.; Hekkelman, M. L.; Hooft, R. W.; Schneider, R.; Sander, C.; Vriend, G. A series of PDB related databases for everyday needs. Nucleic Acids Res. 2010, 39, D411–D419.

(58) McGibbon, R. T.; Beauchamp, K. A.; Harrigan, M. P.; Klein, C.; Swails, J. M.; Herna’ sndez, C. X.; Schwantes, C. R.; Wang, L.-P.; Lane, T. J.; Pande, V. S. MDTraj: a modern open library for the analysis of molecular dynamics trajectories. Biophys. J. 2015, 109, 1528–1532.

(59) Scherer, M. K.; Trendelkamp-Schroer, B.; Paul, F.; P’serez-Hernández, G.; Hoffmann, M.; Plattner, N.; Wehmeyer, C.; Prinz, J.-H.; Noé, F. PyEMMA 2: A software package for estimation, validation, and analysis of Markov models. J. Chem. Theory Comput. 2015, 11, 5525–5542.

(60) Rousseau, R.; Schreiner, E.; Kohlmeyer, A.; Marx, D. Temperature-dependent conformational transitions and hydrogen-bond dynamics of the elastin-like octapeptide GVG (VPGVG): a molecular-dynamics study. Biophys. J. 2004, 86, 1393–1407.

(61) Robustelli, P.; Piana, S.; Shaw, D. E. Developing a molecular dynamics force field for both folded and disordered protein states. Proc. Natl. Acad. Sci. U.S.A. 2018, 115, E4758–E4766.

(62) Weißheit, S.; Kahse, M.; Kämpf, K.; Tietze, A.; Vogel, M.; Winter, R.; Thiele, C. M. Elastin-like Peptide in Confinement: FT-IR and NMR T1 relaxation data. Z. Phys. Chem. 2018, 232, 1239–1261.

(63) Li, B.; Alonso, D. O.; Daggett, V. The molecular basis for the inverse temperature transition of elastin. J. Mol. Biol. 2001, 305, 581–592.

(64) Prinz, J.-H.; Wu, H.; Sarich, M.; Keller, B.; Senne, M.; Held, M.; Chodera, J. D.; Schütte, C.; No’ se, F. Markov models of molecular kinetics: Generation and validation. J. Chem. Phys. 2011, 134, 174105.

(65) Chodera, J. D.; No’ se, F. Markov state models of biomolecular conformational dynamics. Curr. Opin. Struct. Biol. 2014, 25, 135–144.

(66) P’serez-Hernández, G.; Paul, F.; Giorgino, T.; De Fabritiis, G.; Noé, F. Identification of slow molecular order parameters for Markov model construction. J. Chem. Phys. 2013, 139, 07B604 1.

(67) Wehmeyer, C.; Scherer, M. K.; Hempel, T.; Husic, B. E.; Olsson, S.; No’ se, F. Introduction to Markov state modeling with the PyEMMA software [Article v1.0]. LiveCoMS 2019, 1, 5965, DOI: 10.33011/livecoms.1.1.5965.

(68) Röblitz, S.; Weber, M. Fuzzy spectral clustering by PCCA+: application to Markov state models and data classification. Adv. Data Anal. Classif. 2013, 7, 147–179.

(69) De Sancho, D. Phase separation in amino acid mixtures is governed by composition. Biophysical Journal 2022, 121, 4119–4127.

(70) Garanger, E.; MacEwan, S. R.; Sandre, O.; Bruûlet, A.; Bataille, L.; Chilkoti, A.; Lecommandoux, S. Structural evolution of a stimulus-responsive diblock polypeptide micelle by temperature tunable compaction of its core. Macromolecules 2015, 48, 6617–6627.

(71) Bouchaud, J. P. Weak ergodicity breaking and aging in disordered systems. J. Phys. I 1992, 2, 1705–1713, DOI: 10.1051/jp1:1992238.

(72) Rickard, M. M.; Zhang, Y.; Gruebele, M.; Pogorelov, T. V. In-cell protein–protein contacts: Transient interactions in the crowd. J. Phys. Chem. Lett. 2019, 10, 5667– 5673.

(73) Hammouda, B. Polymer Characteristics: SANS from homogeneous polymer mixtures: A unified overview ; Springer Berlin Heidelberg: Berlin, Heidelberg, 1993; pp 87–133, DOI: 10.1007/BFb0025862.

(74) Pesce, F.; Newcombe, E. A.; Seiffert, P.; Tranchant, E. E.; Olsen, J. G.; Kragelund, B. B.; Lindorff-Larsen, K. Assessment of models for calculating the hydrodynamic radius of intrinsically disordered proteins. bioRxiv 2022-06-13, DOI: 10.1101/2022.06.11.495732, (accessed 2022-12-10).

(75) Grimaldo, M.; Roosen-Runge, F.; Zhang, F.; Schreiber, F.; Seydel, T. Dynamics of proteins in solution. Q. Rev. Biophys. 2019, 52, e7, 1–63, DOI: 10.1017/S0033583519000027.

