## Supporting Information for "Structural and Dynamical Properties of Elastin-Like Peptides near their Lower Critical Solution Temperature"

### MSM of a single peptide

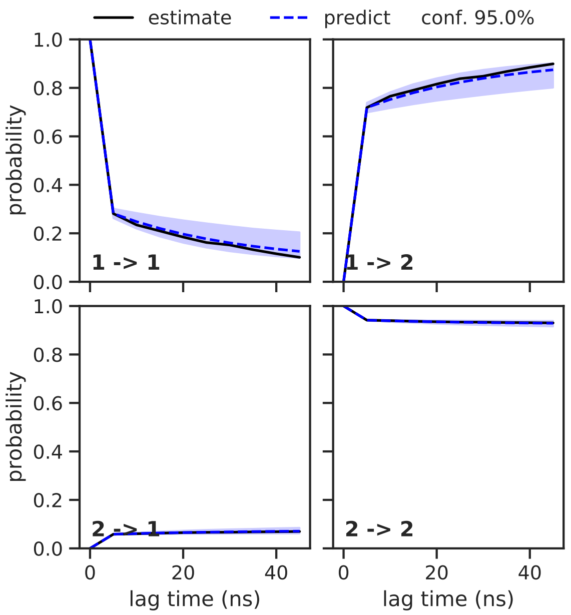


Figure 1: Chapman-Kolmogorov test for the MSM constructed for a GVG(VPGVG)$_3$ peptide using a lag time $\tau$=5 ns and two macrostates. Estimations (blue dashed lines) with confidence interval (blue shades) and predictions (black continuous line) are shown.

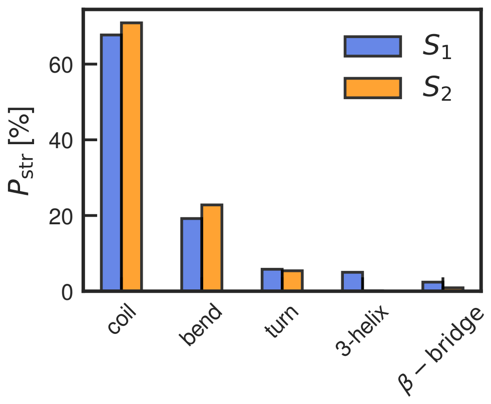


Figure 2: The probability $P_{\text{str}}$ that a residue belongs to a given type of the secondary structure for two MSM states.

### Chain size for sequence-modified peptides

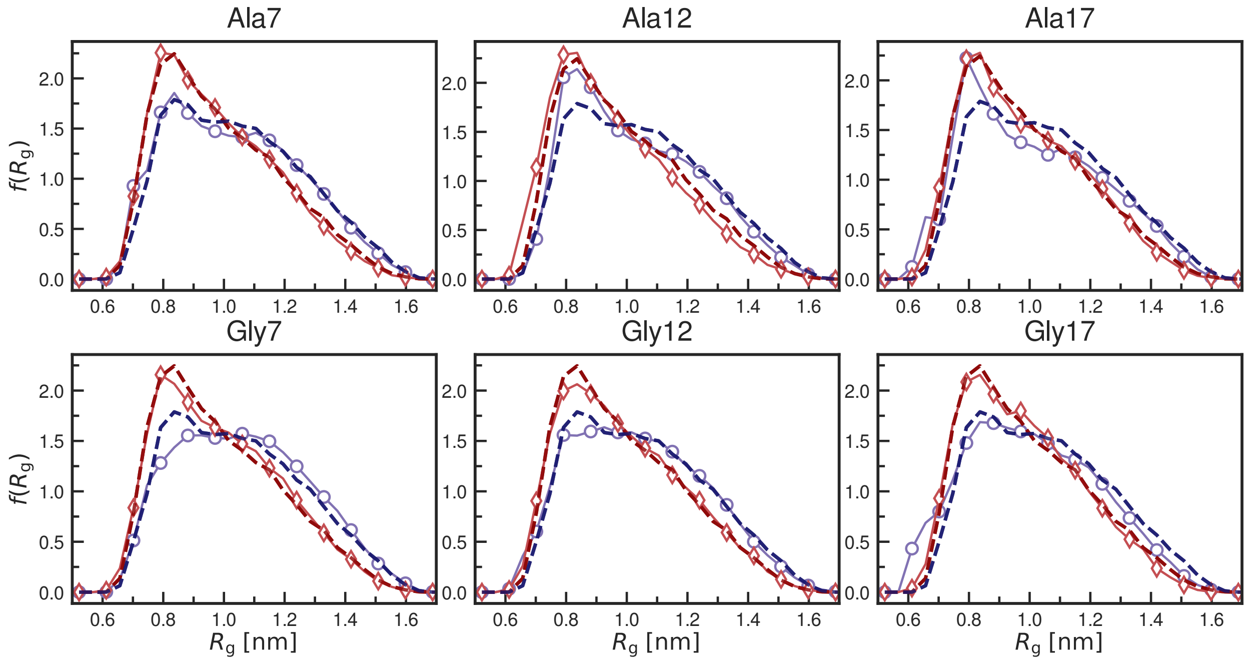


Figure 3: Probability density function of the radius of gyration $f(R_g)$ of a single peptide for six new sequences investigated. Labels indicate the modification made to the original sequence. Dashed dark lines represent the distribution for the original sequence.

### Contact maps for glycine-modified sequences

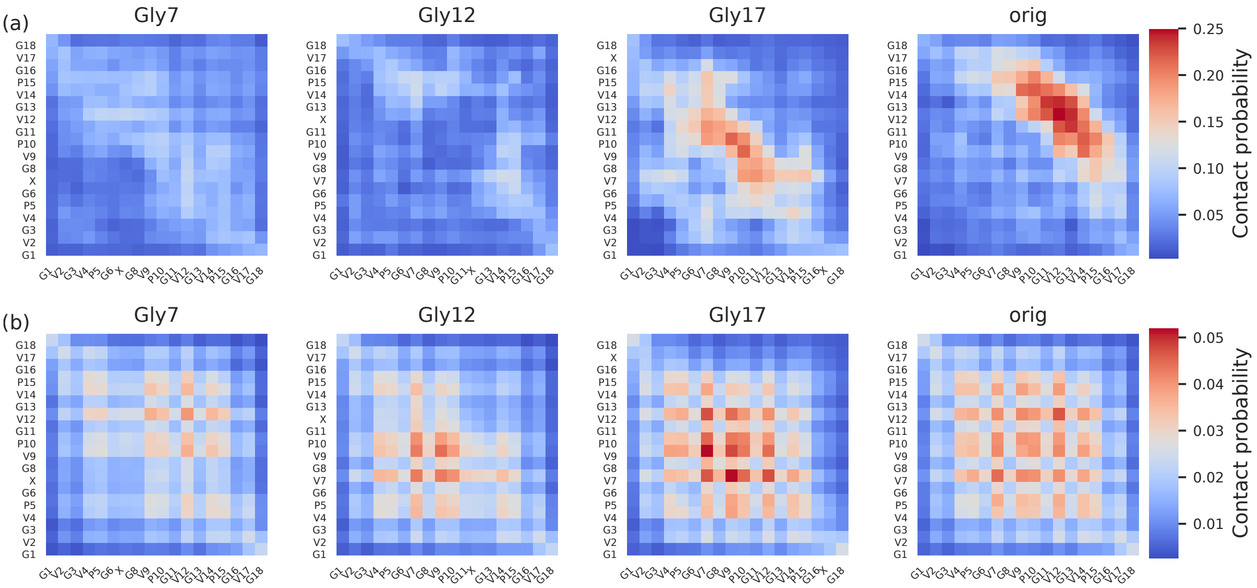


Figure 4: (a) and (b) Interpeptide contact maps for the system at $c_{\mathrm{pept}}$ = 23 mg/ml at temperatures $T$ =288 K and $c_{\mathrm{pept}}$ = 115 mg/ml and $T$ =350 K, respectively. The sequence modification is indicated by a label on top of each contact map. Label "orig" corresponds to the originally studied sequence. A cutoff of 0.5 nm is used to define contact between atoms. Contact maps are constructed using the same scale.

#### Contact lifetime distribution in modified sequences

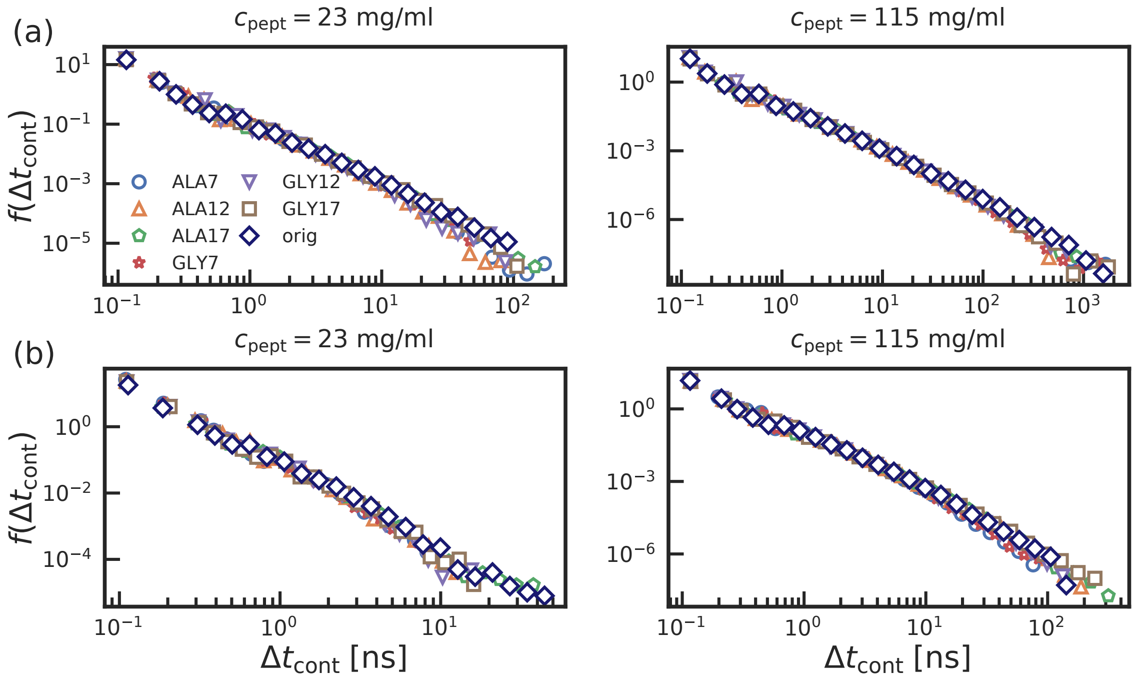


Figure 5: Probability density function of contact time $f(\Delta t_{\mathrm{cont}})$ at temperature (a) T=288 K, and (b) T=350 K for two peptide concentrations investigated.

#### Total number of contacts in multi-peptide systems

In Figure 6, we show the total number of contacts $N_{\mathrm{cont}}^{\mathrm{total}}$ formed during a simulation as a function of temperature across multi-peptide systems investigated.

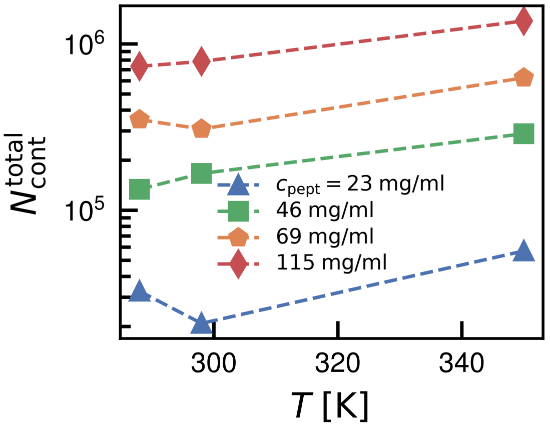


Figure 6: Total number of contacts $N_{\mathrm{cont}}^{\mathrm{total}}$ in multi-peptide systems as a function of temperature $T$.
